# Unraveling Diagnostic Biomarkers of Schizophrenia Through Structure-Revealing Fusion of Multi-Modal Neuroimaging Data

**DOI:** 10.1101/543603

**Authors:** Evrim Acar, Carla Schenker, Yuri Levin-Schwartz, Vince Calhoun, Tülay Adalı

**Affiliations:** Machine Intelligence Department Simula Metropolitan Center for Digital Engineering Oslo, Norway; Department of Environmental Medicine and Public Health Icahn School of Medicine at Mount Sinai New York, New York, United States; The Mind Research Network Albuquerque, New Mexico, United States; Department of Computer Science and Electrical Engineering University of Maryland Baltimore County Baltimore, Maryland, United States

**Keywords:** EEG, fMRI, sMRI, schizophrenia, structural/functional biomarkers, coupled matrix/tensor factorization, ICA

## Abstract

Fusing complementary information from different modalities can lead to the discovery of more accurate diagnostic biomarkers for psychiatric disorders. However, biomarker discovery through data fusion is challenging since it requires extracting interpretable and reproducible patterns from data sets, consisting of shared/unshared patterns and of different orders. For example, multi-channel electroencephalography (EEG) signals from multiple subjects can be represented as a third-order tensor with modes: *subject*, *time*, and *channel*, while functional magnetic resonance imaging (fMRI) data may be in the form of *subject* by *voxel* matrices. Traditional data fusion methods rearrange higher-order tensors, such as EEG, as matrices to use matrix factorization-based approaches. In contrast, fusion methods based on coupled matrix and tensor factorizations (CMTF) exploit the potential multi-way structure of higher-order tensors. The CMTF approach has been shown to capture underlying patterns more accurately without imposing strong constraints on the latent neural patterns, *i.e.*, biomarkers. In this paper, EEG, fMRI and structural MRI (sMRI) data collected during an auditory oddball task (AOD) from a group of subjects consisting of patients with schizophrenia and healthy controls, are arranged as matrices and higher-order tensors coupled along the *subject* mode, and jointly analyzed using structure-revealing CMTF methods (also known as advanced CMTF (ACMTF)) focusing on unique identification of underlying patterns in the presence of shared/unshared patterns. We demonstrate that joint analysis of the EEG tensor and fMRI matrix using ACMTF reveals significant and biologically meaningful components in terms of differentiating between patients with schizophrenia and healthy controls while also providing spatial patterns with high resolution and improving the clustering performance compared to the analysis of only the EEG tensor. We also show that these patterns are reproducible, and study reproducibility for different model parameters. In comparison to the joint independent component analysis (jICA) data fusion approach, ACMTF provides easier interpretation of EEG data by revealing a single summary map of the topography for each component. Furthermore, fusion of sMRI data with EEG and fMRI through an ACMTF model provides structural patterns; however, we also show that when fusing data sets from multiple modalities, hence of very different nature, preprocessing plays a crucial role.

## 1 Introduction

Multiple neuroimaging techniques provide complementary views of neural structure and function. For instance, one of the most commonly used neuroimaging methods, electroencephalography (EEG), measures the electrical activity with high temporal but low spatial resolution, while functional magnetic resonance imaging (fMRI) records the changes in the blood flow with high spatial but low temporal resolution [1, 2]. Therefore, joint analysis of signals from multiple neuroimaging modalities is of interest in order to better understand neural activity and to discover reliable diagnostic biomarkers for psychiatric disorders such as schizophrenia [3, 4, 5, 6, 7].

With the advances in technology, vast amounts of neuroimaging data has been generated; however, data mining or signal processing methods so far have limited success in terms of finding reliable diagnostic imaging biomarkers for many psychiatric disorders [4, 8]. One of the reasons for this limited success has been the fact that data fusion is a particularly challenging task when the goal is to extract reproducible and interpretable patterns. Data from different sources consists of both shared (or common) and unshared (or distinct) underlying patterns [9, 3, 10, 2], and even the definition of “sharedness” is a topic of current research [11, 12]. Furthermore, data sets from different modalities may be of different orders, such as multi-channel EEG signals from multiple subjects can be represented in the form of a third-order tensor with modes: *subject*, *time*, and *channel*, while fMRI data is often represented as a *subject* by *voxel* matrix (Figure 1). Similar challenges have been observed in other disciplines targeting biomarker discovery as well, *e.g.*, in joint analysis of omics data [13], where the ultimate goal is to discover significant metabolites, genes, etc. as potential biomarkers.

**Figure 1:**
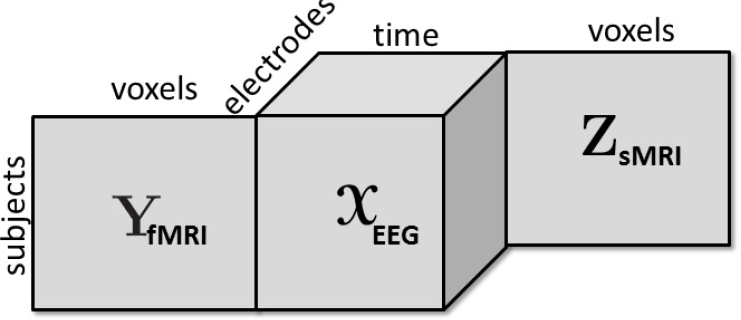
A third-order tensor representing multi-channel EEG signals is coupled with fMRI and sMRI data in the form of matrices in the *subject* mode.

The common approaches for fusion of multi-modal neuroimaging data are based on matrix factorizations, such as joint independent component analysis (jICA) [14], parallel ICA [15] and independent vector analysis (IVA)-based techniques [16], where signals from multiple modalities are represented as matrices, *e.g.*, fMRI data in the form of a *subject* by *voxel* matrix, and EEG signals as a *subject* by *time* matrix [16]. Matrix factorization-based fusion methods require additional constraints to recover patterns uniquely [9, 10, 17, 16] and a common practice in neuroscience is to assume that extracted patterns (i.e., biomarkers, or spatial/temporal patterns) are statistically independent. Drawbacks of the traditional methods are two-fold: (i) in the presence of multi-channel EEG signals, which can naturally be represented as a third-order tensor, data is either matricized in the form of a *subject* by *time-channel* matrix [18] or only the signal from a single channel is analyzed [16], ignoring the potential multilinear structure of multi-channel EEG signals, (ii) statistical independence might be a too strong constraint to impose on the patterns; therefore, methods may fail to capture the true patterns [13].

In contrast, coupled matrix and tensor factorizations (CMTF), introduced more recently, have proven useful in terms of addressing the drawbacks of matrix factorization-based fusion methods by jointly analyzing data sets in the form of matrices and higher-order tensors without imposing constraints on the factors when the higher-order tensors have a defined multilinear structure [13]. CMTF-based approaches factorize higher-order tensors using a tensor factorization model while simultaneously factorizing the data sets in the form of matrices, and enable the exploration of the potential multilinear structure inherent to, for instance, multi-channel EEG signals. Previously, analyzing multi-channel EEG signals using tensor factorizations has shown promising performance in terms of capturing spatial, spectral and temporal signatures of epileptic seizures [19, 20] as well as providing better understanding of brain activity patterns [21, 22, 23], see also [24] for a review. Therefore, recent studies analyzing neuroimaging signals from multiple modalities have arranged multi-channel EEG signals as higher-order tensors and used CMTF-type methods to jointly analyze, *e.g.*, EEG and magnetoencephalography [25, 26], EEG and electro-ocular artifacts [27], and EEG and fMRI [28, 29, 30, 31], or arranged multiple diffusion tensor imaging modalities as a third-order tensor and coupled that with gray matter maps [32]. However, among the CMTF-based methods, the ones assuming that coupled data sets have only shared factors, may fail to capture the underlying patterns in the presence of both shared and unshared factors [33, 34]; therefore, they are not ideal for biomarker discovery.

In this paper, we use a CMTF-based approach to jointly analyze neuroimaging signals from multiple modalities, more specifically, fMRI, sMRI and EEG data, collected during an auditory oddball (AOD) task from a group of subjects consisting of patients with schizophrenia and healthy controls with the goal of unraveling potential diagnostic biomarkers for schizophrenia. To the best of our knowledge, this is the first comprehensive study of a CMTF-based method for biomarker discovery for a psychiatric disorder discussing both strengths and limitations of the proposed framework, building onto our preliminary results in [35, 36]. Furthermore, due to the reproducibility and uniqueness requirements of such an application, we use a structure-revealing CMTF model, known as the advanced CMTF (ACMTF) model [33], to estimate weights of the components in each modality in order to identify shared/unshared factors and quantify the contribution from each modality. Our preliminary studies have shown the promise of the ACMTF model in terms of capturing neural patterns that can differentiate between patients with schizophrenia and healthy controls by jointly analyzing EEG-fMRI signals [36] and EEG-fMRI-sMRI data [35]; however, those two studies used only a subset of electrodes, making it difficult to evaluate the added value of each modality in terms of biomarker discovery. Also, in this paper, we include an additional metric to study the additive value of each modality, and evaluate the performance of the models in terms of clustering subjects from different groups, whereas the previous studies only used the interpretation and statistical significance of extracted patterns in terms of differentiating between groups. Clustering results complement univariate statistical significance tests and show whether combinations of potential biomarkers provide meaningful clusters. We show that EEG analysis using a CP model and joint analysis of EEG, fMRI as well as EEG, fMRI and sMRI reveal statistically significant and biologically meaningful components in terms of differentiating between patients with schizophrenia and healthy controls. In comparison to the results when only the EEG data is analyzed, the incorporation of fMRI signals results in clearer spatial maps and better clustering performance. With the incorporation of sMRI, we obtain structural patterns in addition to temporal and spatial patterns of functional activity without degrading the clustering performance. ACMTF models with different parameter settings have been compared, and based on detailed experiments, we observe that ACMTF consistently reveals similar significant patterns, which provide a concise summary of the topography, while being sensitive to certain parameters for uniqueness.

## 2 Materials and Methods

### 2.1 Background

In this section, we briefly discuss the CP tensor model as well as structure-revealing CMTF and jICA models. Let the third-order tensor 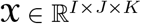 with modes *subject*, *time*, and *electrode*, and matrices **Y** ∈ ℝ^*I*×*M*^ (*subject* by *voxel*) and **Z** ∈ ℝ^*I*×*L*^ (*subject* by *voxel*), represent multi-channel EEG, fMRI and sMRI data, respectively (as in Figure 1).

#### 2.1.1 CANDECOMP/PARAFAC (CP)

The CP model [37, 38], also referred to as the canonical polyadic decomposition [39], is one of the most popular tensor factorization models. It is considered as one of the extensions of the matrix singular value decomposition (SVD) to higher-order tensors (*N* ≥ 3) and represents the tensor as a sum of rank-one tensors. For a third-order tensor 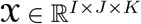, the *R*-component CP model is defined as

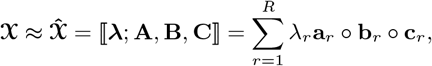

where ∘ indicates the vector outer product. The vectors from the rank-one components are collected in the *factor matrices* **A** ∈ ℝ^*I*×*R*^ = [**a**_1_…**a**_*R*_], **B** ∈ ℝ^*J*×*R*^ = [**b**_1_…**b**_*R*_] and **C** ∈ ℝ^*K*×*R*^ = [**c**_1_…**c**_*R*_]. In this definition, columns of all factor matrices are assumed to be normalized to unit 2-norm and the norms are absorbed in the vector **λ** ∈ ℝ^*R*×1^. For the third-order tensor 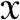 consisting of the EEG data, the factor matrices **A**, **B** and **C** correspond to the extracted factor vectors in the *subject*, *time* and *electrode* modes, respectively. By modeling 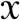 using a CP model, we assume that component *r* models a brain activity with temporal and spatial patterns represented by **b**_*r*_ and **c**_*r*_. Multi-channel EEG signals from each subject are a linear mixture of these *R* brain activities mixed using subject-specific weights. The CP model is also known as a topographic components model [21].

In contrast to matrix factorizations, the CP model of higher order tensors is unique up to scaling and permutation under mild conditions [40, 41], without the need to impose additional constraints.

#### 2.1.2 Structure-Revealing Coupled Matrix and Tensor Factorizations

Given the third-order tensor 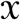 coupled with matrices **Y** and **Z** in the *subject* mode we can jointly factorize them using a structure-revealing CMTF model (*a.k.a.* ACMTF) [33] that fits a CP model to tensor 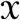 and factorizes matrices **Y** and **Z** in such a way that the factor matrix extracted from the common mode, *i.e.*, *subject*, is the same in the factorizations of all data sets. An *R*-component ACMTF model minimizes the following objective function:

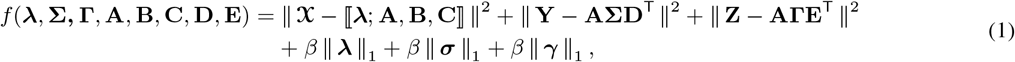

where the columns of factor matrices have unit norm, *i.e.*, || **a**_*r*_ || = || **b**_*r*_ || = || **c**_*r*_ || = || **d**_*r*_ || = || **e**_*r*_ || = 1 for *r* = 1,…, *R*. **λ**, ***σ***, **γ** ∈ ℝ^*R*×1^ are the weights of rank-one terms in 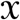, **Y**, and **Z**, respectively. **Σ**, **Γ** ∈ ℝ^*R*×*R*^ are diagonal matrices with entries of ***σ*** and **γ** on the diagonal. **D** ∈ ℝ^*M*×*R*^ and **E** ∈ ℝ^*L*×*R*^ correspond to factor matrices in the *voxel* mode in fMRI and sMRI. ||.|| denotes the Frobenius norm for matrices/higher-order tensors, and the 2-norm for vectors. ||.||_1_ denotes the 1-norm of a vector, *i.e.*, 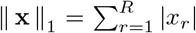 and *β* ≥ 0 is a penalty parameter. Imposing penalties on the weights in (1) sparsifies the weights so that unshared factors have weights close to 0 in some data sets. The model is illustrated in Figure 2. By jointly analyzing neuroimaging data using an ACMTF model, we assume that each component extracted from 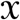 models a brain activity with certain temporal (**b**_*r*_) and spatial (**c**_*r*_) signatures, and the corresponding component in **Y** models related brain activity with higher spatial resolution using **d**_*r*_ while the component in **Z** provides information about the tissue type at a very high spatial resolution using **e**_*r*_. Since the same factor matrix **A** is extracted from the *subject* mode from all data sets, subject-specific coefficients in all modalities are assumed to be the same. The ACMTF model inherits uniqueness from CP [42], as long as all factors are shared, and provides reproducible and interpretable factors. Note that, in the presence of both shared/unshared components, 1-norm penalties on the weights help to obtain unique solutions [33].

**Figure 2:**
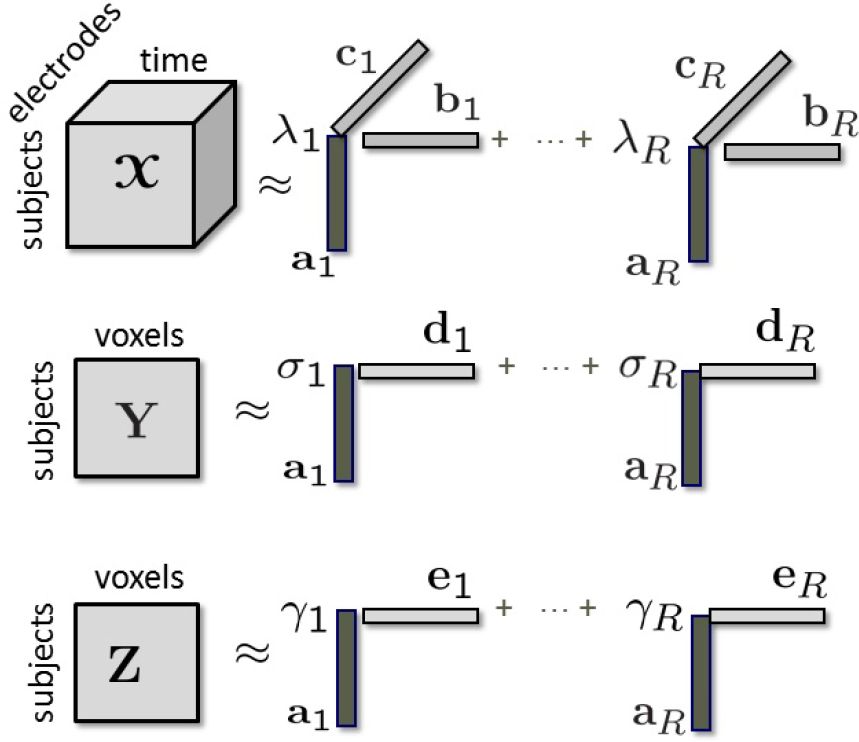
Modeling of tensor 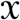 coupled with matrices **Y** and **Z** in the *subject* mode using a structure-revealing CMTF model.

#### 2.1.3 Joint ICA

An alternative approach to jointly analyze 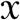, **Y** and **Z** is to use a matrix factorization-based fusion approach by matricizing 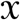 in the *subject* mode as a *subject* by *time - electrode* matrix denoted as **X**_**(1)**_. Joint ICA [14] concatenates matrices representing the data from different modalities and models the constructed matrix using an ICA model as follows:

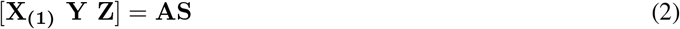

where, for an *R*-component ICA model, **A** ∈ ℝ^*I*×*R*^ corresponds to the mixing matrix, similar to the factor matrix in (1), and **S** ∈ ℝ^*R*×(*JK*+*M*+*L*)^ represents the source signals. Note that subject covariations across all data sets, *i.e.*, modalities, are assumed to be the same in jICA as in ACMTF, since the same mixing matrix is shared across the data sets. However, in this case the model does not include an adaptive estimation of contributions from each modality as in ACMTF, and though this can be captured to a degree within the weights from the estimated components from each modality, it represents a more constrained model. The rows of **S** corresponding to patterns of brain activity are assumed to be statistically independent.

### 2.2 Experiments

We make use of EEG, fMRI, and sMRI data collected from patients with schizophrenia and healthy controls to show the use of ACMTF models to discover potential diagnostic biomarkers for schizophrenia. Our experiments focus on joint analysis of EEG and fMRI data, and discuss the effect of different modeling choices, *i.e.*, number of components (*R*), the penalty parameter (*β*), preprocessing, and use of subsets of electrodes. We also discuss the performance of ACMTF in comparison with jICA. Furthermore, the analysis of only EEG signals and joint analysis of EEG, fMRI, and sMRI have been studied to show the information gain with each modality and potential issues due to the use of additional modalities.

#### 2.2.1 Data

The EEG, fMRI and sMRI data were separately collected from 21 healthy controls and 11 patients with schizophrenia during an auditory oddball task (AOD), where subjects pressed a button when they detected an infrequent target sound within a series of auditory stimuli. For the fMRI data, we computed task-related spatial activity maps for each subject, calculated by the general linear model-based regression approach using the statistical parametric mapping toolbox (SPM 12)^1^. By making use of these features, we constructed a matrix of 32 *subjects* by 60186 *voxels* representing the fMRI signals. For the EEG data, for each electrode, we averaged small windows around the target tone across the repeated instances, deriving event-related potentials. Out of 64 electrodes in total, we used 62 electrodes by excluding the two corresponding to vertical and horizontal electrooculography (EOG) electrodes. Multi-channel EEG signals were then arranged as a third-order tensor: 32 *subjects* by 451 *time samples* by 62 *electrodes*. In order to assess the modeling assumptions, we also used a subset of electrodes from frontal, motor and parietal areas, *i.e.*, AF3, AF4, Fz, T7, C3, Cz, C4, T8, Pz, PO3 and PO, and, in that case, formed a third-order tensor with 11 electrodes as in [35, 36]. For the sMRI data, we computed probabilistically segmented gray matter images for each subject and by using these features formed a matrix of 32 *subjects* by 306640 *voxels*. For more details, see [16].

#### 2.2.2 Experimental Setting

Before the analysis, we centered the third-order EEG tensor across the *time* mode, and scaled within the *subject* mode by dividing each horizontal slice by its standard deviation (see [43] for further details on preprocessing of higher-order tensors). The fMRI and sMRI data were also preprocessed by centering each row (subject-wise) and dividing each row by its standard deviation. When fitting the ACMTF model, each data set was also divided by its Frobenius norm to give equal importance to the approximation of each data set.

In order to demonstrate the information gained by the addition of each modality and sensitivity of the fusion approach to various modeling choices, the following experiments are carried out:

- *Individual analysis of the EEG tensor* using a CP model,
- *Joint analysis of the EEG tensor coupled with fMRI* using an ACMTF model (i) by leaving out signals from one subject at a time, (ii) for 11-electrode vs. 62-electrode case, (iii) in comparison with jICA, (iv) with different number of components, *R*, (v) with different sparsity penalty parameters, *β*, (vi) with/without additional centering across the *subject* mode.
- *Joint analysis of the EEG tensor coupled with fMRI and sMRI* using an ACMTF model.

The CP model is fit using CP-OPT [44] from the Tensor Toolbox version 2.5^2^ using the nonlinear conjugate gradient algorithm (NCG). For the ACMTF model, we use ACMTF-OPT [33] from the CMTF Toolbox, also using NCG to fit the model. Multiple random initializations are used to fit the models, and the solution corresponding to the minimum function value is reported. Furthermore, the ACMTF model is experimentally validated to be unique by obtaining the same minimum function value^3^ a number of times and checking the uniqueness of model parameters, *i.e.*, factor matrices and weights of the factors (up to permutation)^4^. For jICA, we unfold the EEG tensor in the *subject* mode forming a matrix of 32 *subjects* × 27962 (*time-electrodes*), and concatenate the resulting matrix with the fMRI matrix. The concatenated matrix is decomposed using an ICA algorithm based on entropy bound minimization [45], which makes use of a flexible density model that is a better fit to data formed by concatenating signals from different modalities [16]. We fit the model using multiple random initializations and report the most stable run determined by a minimum spanning tree-based approach [46].

### 2.3 Performance Evaluation

The performance is assessed both qualitatively and quantitatively. The qualitative assessment relies on the interpretation of the extracted temporal and spatial patterns as well as comparisons with the previous findings in the literature on schizophrenia. For the quantitative assessment, we perform the following:

- *Two-sample t-test*: Since the ultimate goal of any factorization of this data is the discovery of latent factors that can differentiate between patients with schizophrenia and healthy controls, we can quantify the performance of a method based on its ability to produce factors that can provide such a differentiation. With the assumption of unequal variance for the healthy control and patient groups, a two-sample *t*-test is applied on each column of the factor matrix extracted from the *subject* mode, which is of size 32 by *R*. Out of *R* columns, those that have *subject* mode vectors that are statistically significant, *i.e.*, with p-values < 0.05, are identified and corresponding temporal and spatial patterns are reported as potential biomarkers.
- *Clustering*: Subjects are clustered into two groups based on the factor matrix corresponding to the *subject* mode using *k*-means clustering, where *k*-means is performed 100 times with different initializations and the most consistent cluster assignments are used. Unlike the *t*-test based approach that is performed on each column individually, clustering is performed on all possible combinations of the columns of the factor matrix and the performance of the best combination is reported. Therefore, this approach provides a more global view of the discriminatory power of the resulting factorization than the *t*-test based approach. The clustering performance is assessed in terms of accuracy and *F*_1_-score, where 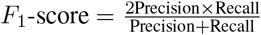, and a patient being clustered as a patient is considered a true-positive.

When assessing different modeling choices, we also report the model fit defined as:

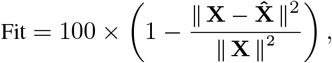

where **X** stands for the raw data (*e.g.*, EEG tensor or fMRI/sMRI matrix), and **X̂** denotes the model. A fit of 100% means that the data is fully explained by the model. The fit shows whether the model explains the data well and indicates the unexplained part left in the residuals. Also, the change in model fit for different number of components shows whether there is a significant gain, in terms of explaining the remaining part in the residuals, by adding more components.

Finally, we compare the similarity of the significant components extracted by different models using a similarity score called the factor match score (FMS). The FMS of component *k* from two models 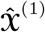 and 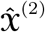 of the EEG tensor 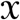 is defined as

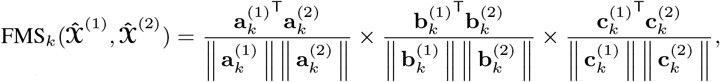

and from two models **Ŷ**^(1)^, **Ŷ**^(2)^ of the fMRI matrix **Y** as

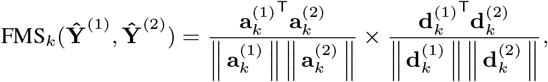

where 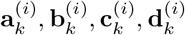 correspond to the *k*th column of the factor matrix corresponding to *subject*, *time*, *electrode* and *voxel* mode of the *i*th model, respectively, after finding the best matching factors for the two models. When components are compared for the models with mismatching dimensions, such as number of subjects or number of electrodes, the mismatching dimension is omitted in the product. An FMS close to one implies similarity of the compared components, while very different components will have an FMS close to zero. FMS is used to quantify the reproducibility of the extracted patterns in addition to qualitative interpretations based on the plots.

For visualization of the extracted components, patterns from fMRI and sMRI *voxel* modes are plotted as z-maps, thresholded at *z* = 2.7, where red indicates an increase in controls over patients and blue indicates an increase in patients over controls. Patterns extracted from the *electrode* mode of the EEG tensor are plotted using the topoplot function from the EEGLAB v13.6.5b [47].

## 3 Results

### 3.1 Individual analysis of the EEG tensor using a CP model

As shown in Figure 3, the CP model of the EEG data in the form of a *subject* by *time* by *electrode* tensor constructed using 62 electrodes has captured significant components in terms of differentiating healthy controls and patients with schizophrenia. The model is fit using *R* = 3 components, and reveals factor matrices **A** ∈ ℝ^32×3^, **B** ∈ ℝ^451×3^, and **C** ∈ ℝ^62×3^ corresponding to *subject*, *time* and *electrode* modes, respectively. The fit of the model is 62% indicating that using only three components a major part of the data can be explained. The *t*-tests performed on the columns of **A** indicate that all three components are significant. The first component, represented in Figure 3 A, corresponds to the third positive peak (P3) and is heavily weighted by central electrodes. The second component, represented in Figure 3 B, refers to the N1-P2 as well as the N2-P3 transitions and is heavily weighted by central and parietal electrodes. The third component, represented in Figure 3 C, refers to the N2 as well as a negative peak after P3 and is heavily weighted by frontal and central electrodes. CP models with different number of components have been fitted to the data as well but those either revealed fewer components with less significance or are degenerate, *i.e.*, a CP model with that many components is not an appropriate model for the data (see [48] for more on degeneracy). Table 1 shows that subjects can be clustered into two groups with 81% accuracy using the factor matrix **A**. Note that clustering performance is similar to the one achieved using the CP model of a tensor constructed using only a subset of electrodes indicating that the assumption of same subject coefficients and temporal patterns across all electrodes is not decreasing the performance. This also may indicate that the additional electrodes are not providing much added information beyond that which is contained by a subset of the electrodes.

**Table 1:** Performance in terms of clustering for different modeling values as well as the factor match scores in comparison to the 10-component ACMTF model (no centering, 62 electrodes).

| | $R$ | Centering | # of Electrodes | Clustering Performance | | FMS | | | |
| --- | --- | --- | --- | --- | --- | --- | --- | --- | --- |
| | | | | Accuracy (%) | $F_1$-score | Component A | | Component B | |
|  |  |  |  |  |  | EEG | fMRI | EEG | fMRI |
| EEG (CP) | 3 | No | 11 | 78 | 0.76 |  |  |  |  |
|  | 3 | No | 62 | 81 | 0.79 |  |  |  |  |
| EEG - fMRI (ACMTF) | 10 | No | 11 | 91 | 0.87 | 0.82 | 0.73 | 0.80 | 0.72 |
|  | 10 | No | 62 | 88 | 0.82 | 1.00 | 1.00 | 1.00 | 1.00 |
|  | 11 | No | 62 | 88 | 0.80 | 0.95 | 0.92 | 0.65 | 0.61 |
|  | 12 | No | 62 | 91 | 0.86 | 0.93 | 0.89 | 0.56 | 0.61 |
|  | 9 | Yes | 62 | 88 | 0.78 | 0.82 | 0.80 | 0.64 | 0.56 |
|  | 10 | Yes | 62 | 91 | 0.88 | 0.85 | 0.81 | 0.57 | 0.57 |
|  | 11 | Yes | 62 | 91 | 0.87 | 0.84 | 0.81 | 0.69 | 0.55 |
|  | 12 | Yes | 62 | 88 | 0.78 | 0.66 | 0.63 | 0.65 | 0.57 |
| EEG - fMRI (JICA) | 10 | No | 62 | 84 | 0.74 |  |  |  |  |
|  | 15 | No | 62 | 82 | 0.70 |  |  |  |  |
|  | 20 | No | 62 | 91 | 0.84 |  |  |  |  |
| EEG - fMRI -sMRI (ACMTF) | 10 | No | 62 | 84 | 0.71 | 0.93 | 0.86 | 0.87 | 0.84 |
|  | 10 | Yes | 62 | 91 | 0.87 | 0.78 | 0.73 | 0.68 | 0.60 |
|  | 15 | Yes | 62 | 91 | 0.86 | - | - | 0.66 | 0.58 |

### 3.2 Joint analysis of EEG and fMRI

Shown in Figure 4, the joint analysis of the EEG tensor and fMRI matrix using an ACMTF model has revealed significant components in terms of differentiating between healthy controls and patients while also providing spatial patterns in much higher resolution and improving the clustering performance compared with the CP model of the EEG tensor. The 10-component ACMTF model extracts factor matrices **A** ∈ ℝ^32×10^, **B** ∈ ℝ^451×10^, **C** ∈ ℝ^62×10^, and **D** ∈ ℝ^60186×10^ corresponding to *subject*, *time*, *electrode*, and *voxel* modes, respectively, as well as weights of the components in EEG (λ ∈ ℝ^10×1^) and fMRI (*σ* ∈ ℝ^10×1^). The sparsity penalty parameter is set to *β* = 10^−3^. The fit is 79% and 65% for EEG and fMRI, respectively, indicating that the extracted factors, which have high weights in both data sets indicating shared factors, account for a large part of both data sets. The *t*-test on the columns of **A** reveals that out of ten components, only two, components 1 and 9, are statistically significant. Figure 4 A and B illustrate the two significant components in *time* (**b**_*r*_), *electrode* (**c**_*r*_) and *voxel* (**d**_*r*_) modes. Figure 4 C shows component weights in each data set. From Figure 4, we see that though both significant components have a contribution from both EEG and fMRI, the contribution from EEG to each of these components is greater. This means that the discriminatory information plays a larger part in EEG relative to the other components in EEG than it does for the fMRI. The first component, shown in Figure 4 A, is similar to the component shown in Figure 3 C, and refers to the P2-N2 transition as well as the P3 peak and is heavily weighted by the frontal and central electrodes. A similar activation pattern is seen in the positive activations in the fMRI, though with greater spatial resolution. The second component, shown in Figure 4 B, shares some similarity with the component shown in Figure 3 A, since both are related to the P3 peak but this component is more heavily weighted by the frontal electrodes. The fMRI in Figure 4 B indicates a decrease in activation of controls versus patients in parts of the anterior sensorimotor cortex. We should note that there are some similarities between the areas highlighted in the topographic maps and the regions highlighted in the fMRI. The areas of increased activation of controls over patients in the fMRI, namely frontal and sensorimotor, generally correspond to the greatest weights in the topographic maps. Similar components have been found previously in other analyses of similar data [36], thus increasing our confidence in the results. In comparison to the individual analysis of the EEG tensor, the clustering performance is also much higher, *i.e.*, 88% accuracy and *F*_1_ score of 0.82, as shown in Table 1. This indicates that the ACMTF reveals more discriminatory factors through the inclusion of complementary information from the fMRI.

**Figure 3:**
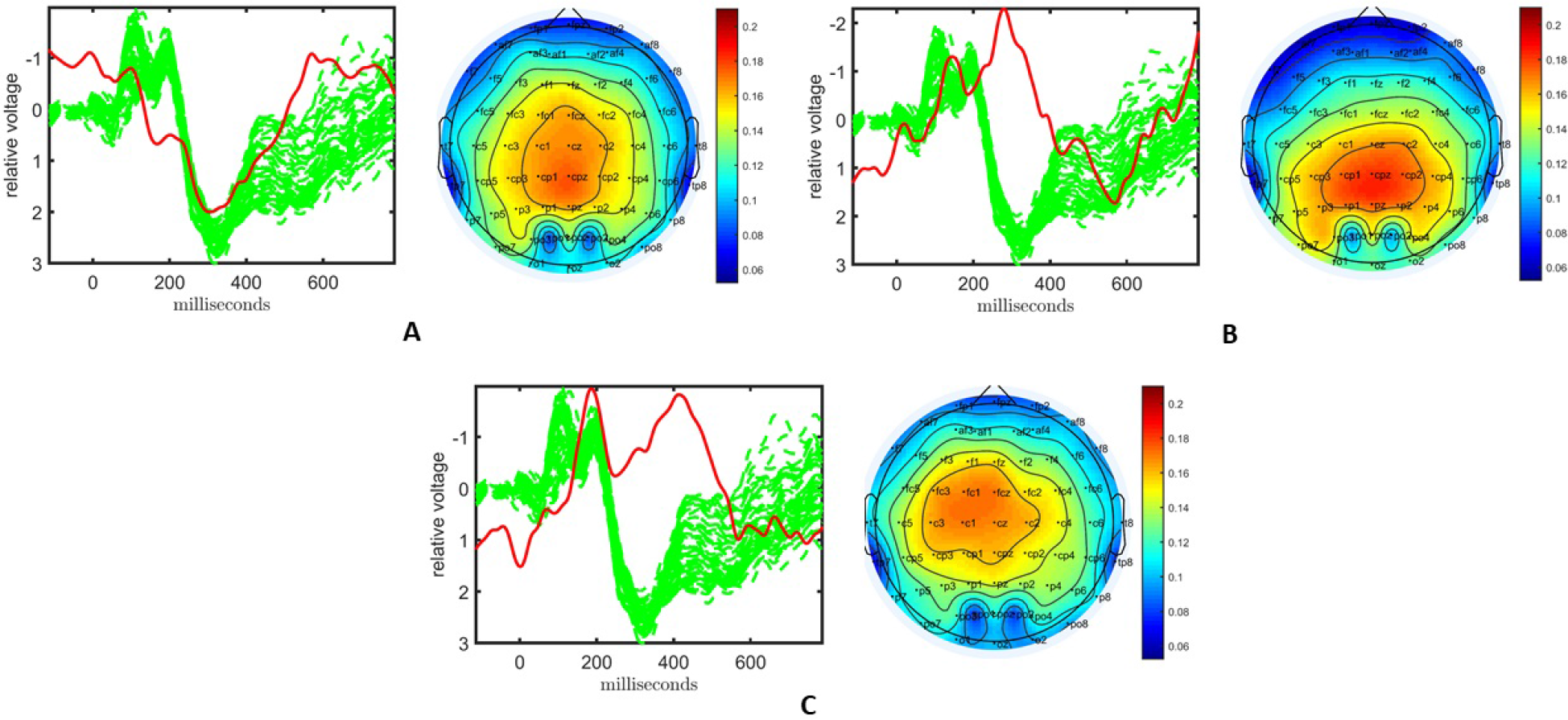
Temporal and spatial patterns from the statistically significant components of the CP model. **(A)** Component 1 corresponds to the P3 peak mainly represented by central electrodes, **(B)** Component 2 refers to the N1-P2 as well as N2-P3 transitions, with high contributions from central and parietal electrodes, **(C)** Component 3 refers to the N2 as well as a negative peak after P3, heavily weighted by frontal and central electrodes. The corresponding *p*-values are 2.1 × 10^−3^, 1.6 × 10^−2^, 1.4 × 10^−4^, respectively. Columns of the factor matrix in the *time* mode are in red while green plots show signals from individual electrodes averaged across all subjects.

#### 3.2.1 Sensitivity Analyses

##### Leave-one-out

The patterns captured in different modes using an ACMTF model are reproducible in case of changes in data sets. In order to evaluate the consistency of the results to changes in the original data, we leave out one subject at a time and fit the ACMTF model using the same parameters (*i.e.*, *R* = 10, *β* = 10^−3^). In other words, we construct 32 different EEG-fMRI data set pairs (with 31 subjects) and compare the significant factors extracted using the ACMTF model of each pair with the model derived using 32 subjects. Table 2 shows that average FMS for component 1 (Figure 4 A), which is the most significant factor, is 0.98 for EEG and 0.95 for fMRI indicating close to exact recovery of the same patterns. Average FMS for the less significant component, *i.e.*, component 9 (Figure 4 B), is around 0.90 indicating similar patterns. Furthermore, the average clustering performance is the same as the performance of the original model estimated using data from 32 subjects.

**Table 2:** Leave-one-out sensitivity analysis: Average values (standard deviation) of FMS, clustering performance and fit of the models built on data sets with 31 subjects.

| FMS |  |  |  | Clustering Performance |  | Fit (%) |  |
| --- | --- | --- | --- | --- | --- | --- | --- |
| Component A | | Component B | | Accuracy (%) | $F_1$-score | EEG | fMRI |
| EEG | fMRI | EEG | fMRI |  |  |  |  |
| 0.98 (0.05) | 0.95 (0.09) | 0.92 (0.16) | 0.90 (0.16) | 87.7 (2.4) | 0.81 (0.04) | 79.5 (0.4) | 65.6 (0.3) |

##### 11 electrodes vs. 62 electrodes

When jointly analyzing EEG and fMRI, ACMTF achieves a slightly better performance using a subset of electrodes from certain regions of interest during the construction of the EEG tensor than the case where all 62 electrodes are used to construct the tensor. In our preliminary studies [36], we observed similar components when comparing the 11-electrode case with 62-electrode case while achieving higher statistical significance and better interpretability using 11 electrodes. These observations are also supported by our findings in this study on a slightly different set of subjects (38 subjects in [36] vs. 32 subjects in this paper). Figure 5 illustrates the two most significant components captured by a 10-component ACMTF model of the EEG tensor with 11 electrodes and fMRI data^5^. The fit is 80% and 65% for EEG and fMRI, respectively. As previously observed, both components have higher significance thus indicating that the additional electrodes are not contributing much additional discriminatory information compared with the original 11 electrodes. This may also be related to the fact that the most contributing electrodes to the components in Figure 4 are the electrodes that are part of the set of 11 electrodes. The first component, shown in Figure 5 A, is similar to the component shown in Figure 4 A and refers to P2-N2 transition as well as the P3 peak though with more parietal activation in the fMRI. The second component, shown in Figure 5 B, is similar to the component shown in Figure 4 B and refers to the P3 peak though has more parietal activation in the fMRI, similar to the default mode network. When significant components from 11- and 62-electrode cases are compared, FMS is 0.82 for EEG (excluding the electrode mode) and 0.73 for fMRI for the component given in Figure 4 A, and 0.80 for EEG and 0.73 for fMRI for Figure 4 B. These scores indicate that components are similar to some extent but are not identical. Table 1 indicates slightly higher clustering performance for the 11-electrode case. To summarize, considering that there is minimal difference in performance beyond a slight increase in significance, and that the factors are similar, using all electrodes is preferable to choosing a subset of electrodes, as the latter requires prior knowledge about the functionally relevant electrodes to select and may also introduce a bias by targeting specific regions.

**Figure 4:**
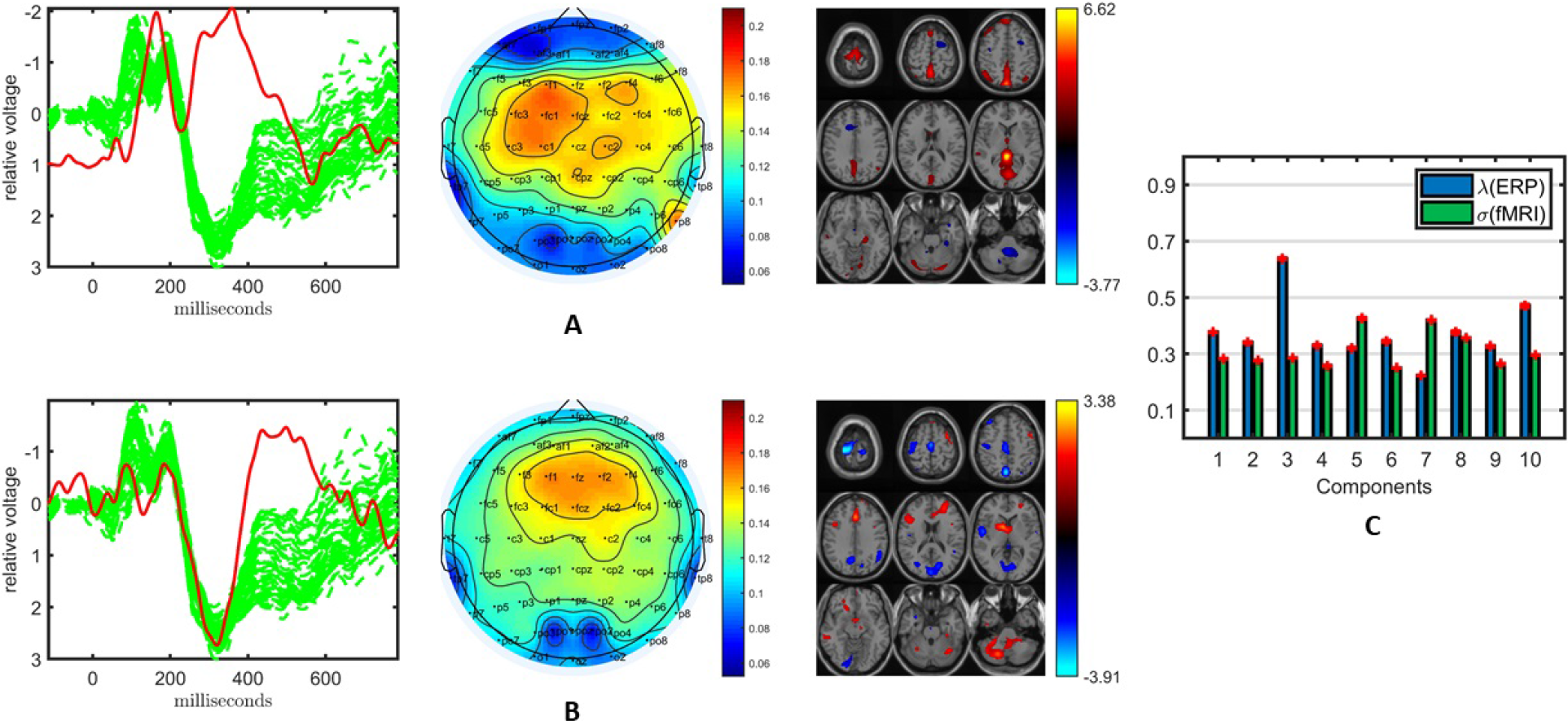
Temporal and spatial patterns from the statistically significant components of the ACMTF model of the EEG tensor with 62 electrodes and fMRI data, with *R* = 10. **(A)** Component 1 refers to the P2-N2 transition as well as the P3 peak, heavily weighted by the frontal and central electrodes in the EEG, and the fMRI shows increased activation of controls over patients in the sensorimotor cortex, **(B)** Component 9 is related to the P3 peak, heavily weighted by the frontal electrodes in the EEG and the fMRI indicates a decrease in activation of controls versus patients in some regions of the sensorimotor cortex and parietal lobe, **(C)** Weights of the components in each data set. The corresponding *p*-values are 6.2 × 10^−3^, 1.9 × 10^−2^, respectively. Columns of the factor matrix in the *time* mode are in red while green plots show signals from individual electrodes averaged across all subjects.

##### ACMTF vs. JICA

The traditional fusion approach jICA can also capture components that can differentiate between healthy controls and patients; however, jICA provides less interpretable patterns. For jICA, the EEG tensor unfolded in the *subject* mode is concatenated with the fMRI data resulting in a 32 (*subject*) by 88148 (*time* × *electrode* - *voxel*) matrix. When this matrix is modeled using jICA with *R* = 10, 15, 20 components, the 10-component model reveals a single component that may be considered statistically significant but the *p*-value is 0.05. The 15-component model reveals a more significant component as illustrated in Figure 6. JICA captures neither a single temporal pattern for all electrodes nor a spatial pattern for all time points, making the intepretation of the components more difficult. In order to provide insight into the topology, spatial patterns as in ACMTF can be computed post hoc from the analysis, e.g., by using peak value of each channel to construct a spatial map for each component [49]; however, that comes with additional assumptions and does not reveal the underlying patterns as naturally as a tensor factorization-based approach. The component in Figure 6 is related to the P2-N2 and N2-P3 transition and the fMRI map includes the expected temporal lobe and default mode regions. We should note that using a 20-component model, a component similar to the one in Figure 6 is captured, indicating that jICA has some stability in regards to the value of *R*. Table 1 shows that while the clustering performance of jICA is lower compared with ACMTF models of EEG and fMRI data sets for *R* = 10 and *R* = 15, it is similar for *R* = 20, indicating that methods with different assumptions may perform the best with different number of components.

**Figure 5:**
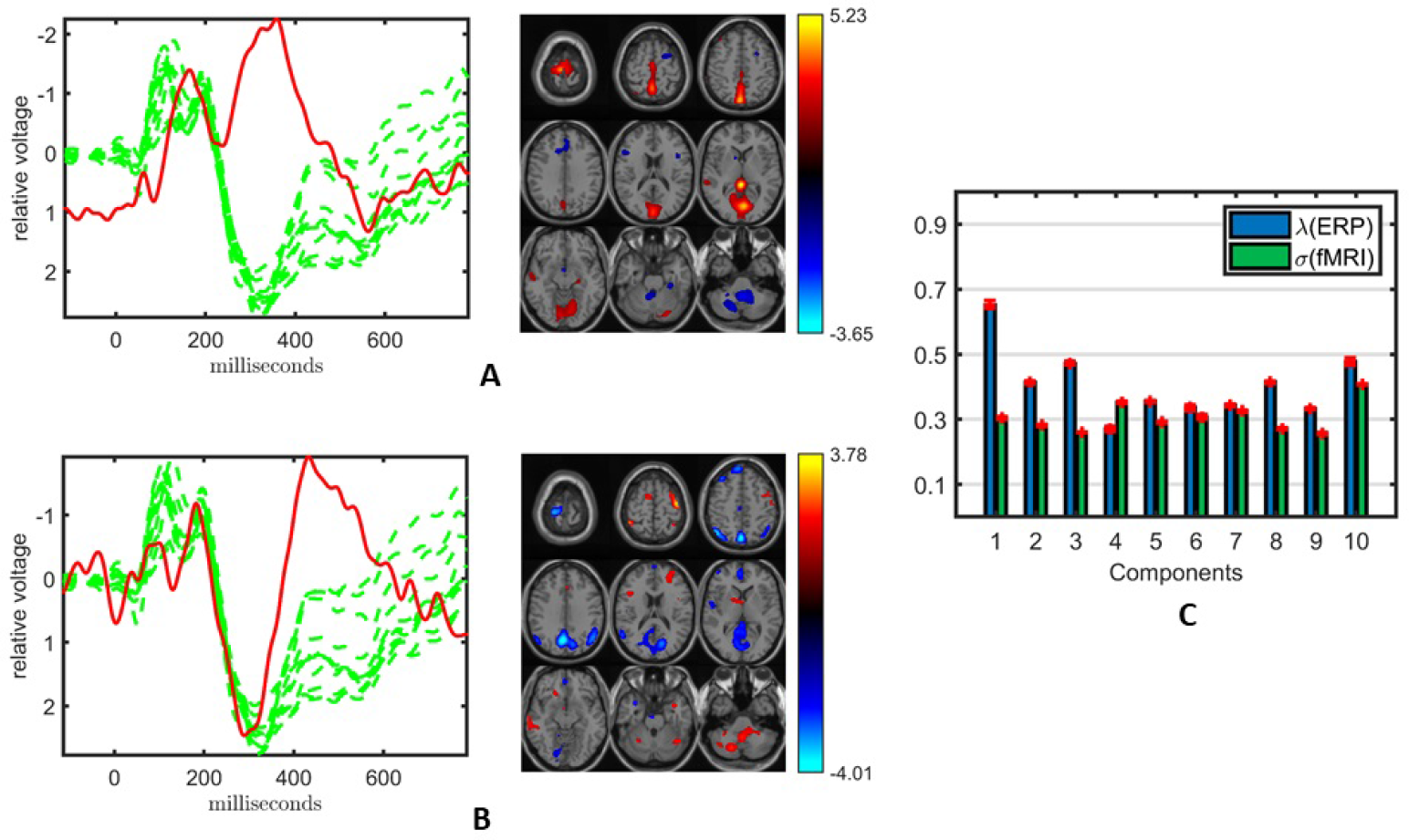
Temporal and spatial patterns from the statistically significant components of the ACMTF model of the EEG tensor with 11 electrodes and fMRI data, with *R* = 10. **(A)** Component 10 corresponds to the P2-N2 transition as well as the P3 peak in the EEG with an increase in sensorimotor and parietal activation of controls over patients in the fMRI, **(B)** Component 5 refers to the P3 peak with a decrease in default mode activity of controls versus patients in the fMRI, **(C)** Weights of the components in each data set. The corresponding *p*-values are 4.3 × 10^−3^, 8.0 × 10^−3^, respectively. Columns of the factor matrix in the *time* mode are in red while green plots show signals from individual electrodes averaged across all subjects.

**Figure 6:**
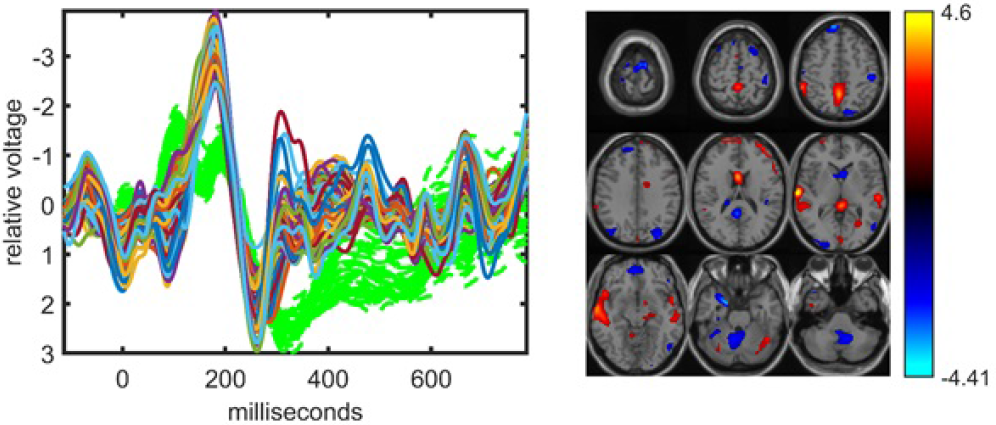
The statistically significant component captured by the jICA model fitted to the concatenation of the unfolded EEG tensor (with 62 electrodes) and fMRI, with *R* = 15. The corresponding *p*-value is 5.2 × 10^−3^. Parts of **s**_*r*_ corresponding to the *time samples* for each electrode in EEG and *voxels* in fMRI are plotted. The EEG part is related to the P2-N2 and N2-P3 transitions. The fMRI indicates some increased activation in the temporal lobe of controls versus patients as well as some posterior cingulate representing the default mode network. In the EEG plot, green dashed plots show signals from individual electrodes averaged across all subjects.

##### Parameter Selection

The ACMTF model is sensitive to two parameters, *i.e.*, the number of components, *R*, and the sparsity penalty parameter, *β*. So far, *R* and *β* are set to *R* = 10 and *β* = 10^−3^. In order to probe the effect of the model order, *R*, we have increased the number of components until the model fails to give a unique solution. We find that as we increase the number of components, the most significant component (*i.e.*, Figure 4 A) is still consistently captured; however, with a decreasing level of similarity. With both *R* = 11 and *R* = 12, the ACMTF model is still unique and reveals significant components that can differentiate between patients and healthy controls. The fit is 81% for EEG and 68% for fMRI with *R* = 11 while 82% for EEG and 70% for fMRI with *R* = 12, indicating that additional components do not explain much additional information. Table 1 shows that a component with a FMS score around 0.90 when compared with the most significant component in a 10-component model (Figure 4 A) has been revealed by both models. The less significant component (Figure 4 B), on the other hand, has limited similarity of around FMS 0.60 with the captured components. The clustering performance for both *R* = 11 and *R* = 12 is similar to that of *R* = 10. Figure S1 (in Supp. Material) illustrates the significant components captured by the ACMTF model with *R* = 12 components. When *R* is increased any further, we cannot obtain a unique solution.

The sensitivity of an ACMTF model to different values of *β* has been studied in [33] using simulated data sets, and it has been shown that in the presence of both shared and unshared components, small values such as *β* = 10^−3^ or *β* = 10^−4^ are effective in terms of uniquely recovering the underlying patterns. For *β* = 0, which corresponds to a CMTF model, the model fails to give a unique solution. For larger values, such as *β* = 10^−2^, it is still possible to find the true solution but the algorithm is very sensitive to the initialization. When EEG and fMRI data sets are jointly analyzed using a 10-component model, weights of the components shown in Figure 4 C indicate that all components are shared. However, even in the presence of only shared components, *β* = 10^−4^ fails to give a unique solution when we get the same function values for different runs while for *β* = 10^−2^, it is not possible to reach to the same function values even with many random initializations due to the sensitivity to initialization.

##### Preprocessing

In addition to the preprocessing steps described in Section 2, in data fusion studies, further preprocessing may be needed, in particular, when the average behavior across subjects accounts for a large variation in one of the data sets vs. the other. SMRI is such a data set and we perform additional centering when we include sMRI in the analysis. Here, when we only consider the joint analysis of EEG and fMRI, an additional centering step across the subjects mode does not affect the clustering performance of the ACMTF model and the significant component in Figure 4 A has also been captured with FMS between 0.80-0.85 (for different number of components *R* = 9, 10, 11) as shown in Table 1. FMS drops for *R* = 12. The less significant component (Figure 4 B), on the other hand, is also estimated, but with FMS within the range 0.55-0.69. Figure S2 (in Supp. Material) illustrates the significant components captured by an ACMTF model with *R* = 12 components. It is important to note that in this case, despite the low FMS values, temporal and spatial patterns are similar to the ones observed in Figure 4 and the interpretation of these two components is the same.

### 3.3 Joint analysis of EEG, fMRI and sMRI

Inclusion of the sMRI data introduces several issues highlighting challenges in data fusion, in particular, preprocessing. If a joint analysis of EEG, fMRI and sMRI data is carried out using an ACMTF model after the preprocessing steps described in Section 2.2.2, the model has two statistically significant components in terms of differentiating healthy controls and patients. Figure 7 illustrates the temporal patterns as well as functional/structural spatial patterns revealed by the significant components. Both components are similar to the components shown in Figure 4 with FMS values between 0.84 - 0.93 as given in Table 1 as well as components shown in Figure 3. However, now information from the two functional modalities, EEG and fMRI, has been combined with information from the structural modality, sMRI. In Figure 7 A, the sMRI portion of the component shows an increase in concentration of gray matter in controls over patients in sections of the parietal lobe and cerebellum. In Figure 7 B, the sMRI portion of the component shows increases in gray matter for controls over patients in multiple portions of the frontal and parietal lobes. Overall these components are similar to components found previously in other analyses of data from the same subjects but with 11 electrodes [35].

Weights of the components, shown in Figure 8, indicate that sMRI does not contribute much to the significant components. In most components the weights of the components in the sMRI are low, indicating potentially unshared factors. However, a closer look at the model reveals that the model fit is 97% for sMRI, while it is 79% for EEG and 64% for fMRI (see Figure S3 in Supp. Material for the singular value spectrum of each data set), and components with high weights in sMRI are mainly modeling the average structure across subjects with highly correlated components, *i.e.*, the correlation of component vector in the *voxel* mode is 0.95 for components 1 and 5. Therefore, data sets must be centered across the *subject* mode to incorporate information other than the mean from sMRI into the analysis. Furthermore, as a result of including components from sMRI, which do not differentiate between the groups, we also observe a drop in the clustering performance in Table 1. However, we do note that the significance of both components have increased, thus indicating that there is additional discriminatory information that the sMRI is providing.

**Figure 7:**
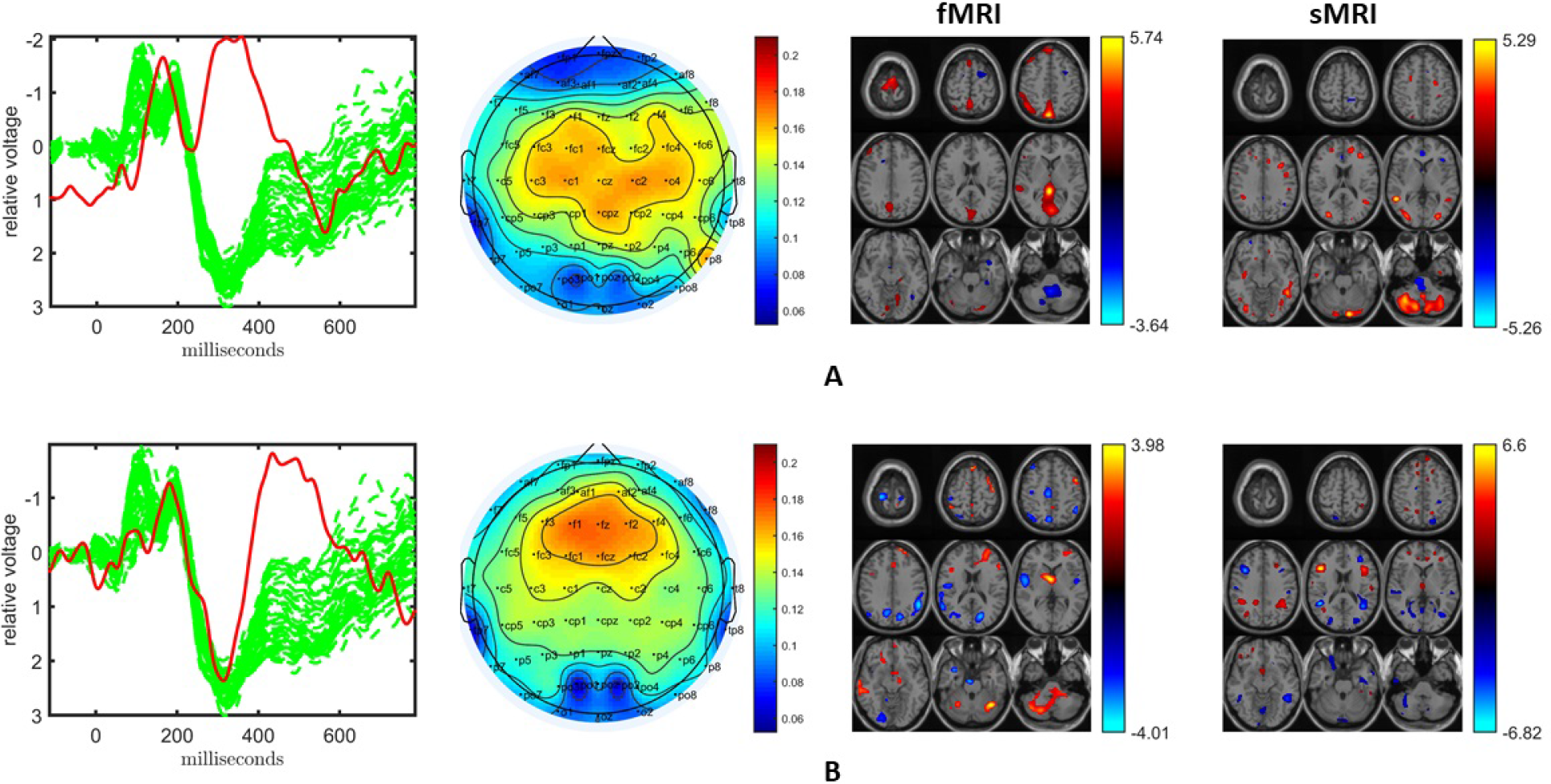
Temporal and spatial patterns from the statistically significant components of the ACMTF model of the EEG tensor with 62 electrodes, fMRI and sMRI data, with *R* = 10. **(A)** Component 10 refers to the P2-N2 transition as well as the P3 peak, heavily weighted by the frontal and central electrodes in the EEG, an increase in sensorimotor and parietal activation of controls over patients in the fMRI, and an increase in concentration of gray matter in controls over patients in sections of the parietal lobe and cerebellum in the sMRI, **(B)** Component 6 corresponds to the P3 peak heavily weighted the frontal electrodes in the EEG, the fMRI indicates a decrease in activation of controls versus patients in some regions of the sensorimotor cortex and parietal lobe, and increases in gray matter for controls over patients in multiple portions of the frontal and parietal lobes in the sMRI. The corresponding *p*-values are 8.0 × 10^−3^, 4.0 × 10^−3^, respectively. Columns of the factor matrix in the *time* mode are in red while green plots show signals from individual electrodes averaged across all subjects.

**Figure 8:**
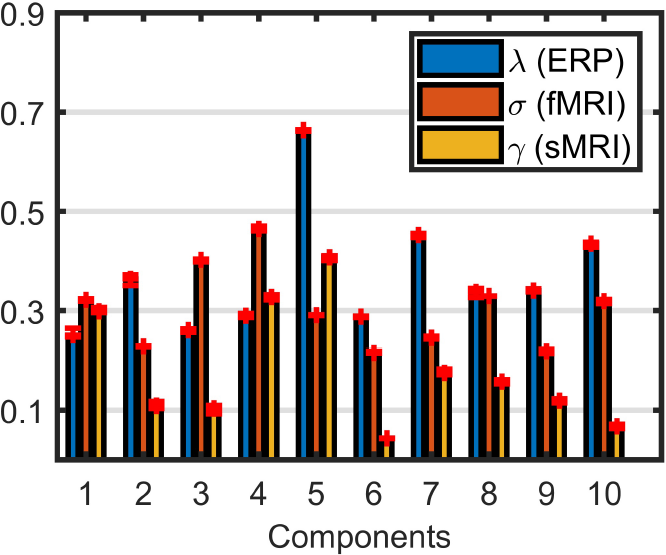
Weights of the components in EEG, fMRI and sMRI extracted by the ACMTF model of the EEG tensor with 62 electrodes, fMRI and sMRI data, with *R* = 10 and no additional centering across the *subject* mode.

When all data sets are centered across the *subject* mode, the ACMTF model has three statistically significant components, which are illustrated in Figure S4 (in Supp. Material). The two most significant ones, shown in Figure S4 A and B, are similar to the components in Figure 4, also indicated by the FMS values in Table 1. The third component represents the N2 peak as well as the P3 peak in the EEG. The topographic map indicates activation in the parietal lobe, while the fMRI shows increased activation of controls over patients in the sensorimotor cortex and a decrease in activation of controls versus patients in the occipital lobe. The sMRI portion of the component indicates changes to gray matter concentration throughout the frontal and parietal lobes. Note that the clustering performance of the overall model has improved significantly by modeling more relevant structure in sMRI compared to the case where there is no centering. The model fit is 69%, 47%, and 70% for EEG, fMRI and sMRI, respectively. In order to increase the model fits, in particular for fMRI, when we increase the number of components to *R* = 15, a unique model can still be obtained with model fits 78%, 64% and 79% for EEG, fMRI and sMRI, respectively. In that case, however, only a single statistically significant component (*p*-value =1.4 × 10^−4^) that is similar to Figure 4 B, is captured. The clustering performance of the 15-component ACMTF model is similar to the 10-component case. These observations indicate that the model is consistent to some degree across models with different numbers of components, by still revealing one of the significant components. The additional components, which explain some of the remaining information in the data sets, do not explicitly contribute to additional significant components but may help with increasing the significance of the relevant component - without hurting the overall clustering performance.

## 4 Discussion

In this paper, we have jointly analyzed multi-modal neuroimaging signals, *i.e.*, EEG, fMRI and sMRI, collected from patients with schizophrenia and healthy controls, using a structure-revealing CMTF model. The model captures temporal as well as functional/structural spatial patterns that can differentiate between patients and healthy controls. Compared to traditional fusion approaches such as jICA, the structure-revealing CMTF model enables us to exploit the multilinear structure of multi-channel EEG signals providing both interpretable patterns and improved uniqueness properties without imposing additional constraints on the extracted patterns. Through joint analysis of EEG, and fMRI, the following temporal and spatial patterns are identified as potential biomarkers:

- *Pattern 1:* The *temporal* part referring to the P2-N2 transition as well as the P3 peak, and the *functional spatial* part showing increased activation of controls over patients in the sensorimotor cortex.
- *Pattern 2:* The *temporal* part referring to the P3 peak, and the *functional spatial* part indicating a decrease in activation of controls versus patients in some regions of the sensorimotor cortex and parietal lobe.

The biomarkers that are extracted using the ACMTF model correspond to signals observed in previous investigations of the structural and functional impacts of schizophrenia. The EEG signals are similar to those observed in previous schizophrenia research [50, 51]. Additionally, the regions in spatial patterns have also been shown to be affected in patients with schizophrenia previously [52, 53, 54]. Through the incorporation of the sMRI data, these patterns have been complemented with the following *structural spatial* parts: (i) *Pattern 1*, the *structural spatial* part indicating an increase in concentration of gray matter in controls over patients in sections of the parietal lobe and cerebellum. (ii) *Pattern 2*, the *structural spatial* part showing increases in gray matter for controls over patients in multiple portions of the frontal and parietal lobes. All three regions have been shown to be impacted in patients with schizophrenia [55, 56, 57, 58]. These patterns are reproducible and have been revealed even in the case of changes in data sets, as we have illustrated by leaving out data from one subject at a time and in our preliminary studies on a slightly different set of subjects [36] and using a subset of electrodes [35].

Any method targeting biomarker discovery must capture the underlying patterns corresponding to the potential biomarkers uniquely; therefore, in this paper, we have used the structure-revealing CMTF model that focuses on unique identification of underlying patterns when jointly analyzing multi-modal data sets with shared and unshared factors, rather than other CMTF methods that have proved useful in missing data estimation applications (where uniqueness of underlying patterns is not of interest) [59, 60]. In addition to the patterns interpreted as potential biomarkers, the structure-revealing CMTF model also reveals weights of the patterns that can be used to identify shared/unshared patterns in each data set and quantify the contribution from each data set. In joint analysis of EEG and fMRI, all components including the statistically significant components differentiating between patients and controls, correspond to shared components. Similarly, in joint analysis of EEG, fMRI and sMRI, as long as necessary preprocessing steps such as centering are carried out as discussed in Section 3.3, all components are shared among three modalities as shown in Figure S5 (in Supp. Material).

One potential drawback of the structure-revealing CMTF model, on the other hand, is sensitivity to its parameters, *i.e.*, the number of components, *R*, and the sparsity penalty parameter, *β*. Despite the sensitivity, we have consistently observed similar statistically significant patterns for different number of components, as shown in Table 1 in terms of FMS values. Note that FMS takes into account every entry in the factor vectors (*i.e.*, many voxels in the fMRI) and it is rather a strict measure. Therefore, we have observed that even for lower FMS values, interpretations of the captured patterns are the same visually (e.g., Figure 4 vs. Figure S2). Another challenge as a result of sensitivity to model parameters is that the model must be experimentally validated to be unique. An important future research direction is to study the landscape of the optimization problem and develop ways to make the problem less sensitive to parameters as well as the initialization. Furthermore, while we have used the same sparsity penalty parameter for all data sets in this paper, in some applications, this parameter may need to be data-specific depending on the structure of each data set.

While data fusion methods are of interest in many disciplines, preprocessing steps have not been well studied within the framework of data fusion. In this paper, we have shown that while centering across the *subject* mode does not affect the joint analysis of EEG and fMRI data, it has a dramatic effect when the sMRI data is incorporated, and the interpretation of the component weights changes significantly. In addition to such preprocessing steps, there are further steps that should be carefully incorporated to data fusion methods such as outlier removal, residual analysis, which may also enable better tools for selecting the number of components.

This paper is a systematic study of the structure-revealing CMTF model for biomarker discovery but with limited number of subjects. In order to see the real promise of the method as a biomarker discovery approach and assess the validity of the potential biomarkers for schizophrenia, joint analysis of EEG, fMRI and sMRI signals must be carried out on a larger set of subjects and also including patients with different neurological disorders. Understanding the promise and limitations of such CMTF-based approaches is crucial as there are increasingly more studies exploiting the multilinear structure of different neuroimaging signals [61, 62, 63, 64, 65, 66, 31, 67].

## Ethics Statement

The protocol was reviewed and approved by the IRB at Hartford Hospital and all participants provided written, informed, consent.

## Author Contributions

EA and TA conceived the project. EA, YLS, and TA designed the experiments. EA and CS performed the data analysis. YLS, TA, and VC interpreted the extracted patterns. EA, CS, YLS, and TA were involved in the preparation of the manuscript. All authors have given approval to the final version of the manuscript.

## Supplemental Data

Supplementary Material has been included.

## Footnotes

1 https://www.fil.ion.ucl.ac.uk/spm/

2 http://www.sandia.gov/tgkolda/TensorToolbox/

3 up to the sixth decimal place

4 Depending on the difficulty of the problem, 160-200 random initializations are used to check for uniqueness.

5 T-tests reveal, in total, four statistically components for this model. However, the other two components, not illustrated in Figure 5, are less significant and have lower FMS values.

