## Supplemental Material for "Unraveling Diagnostic Biomarkers of Schizophrenia Through Structure-Revealing Fusion of Multi-Modal Neuroimaging Data"

A Preprint

### Supplementary Tables and Figures

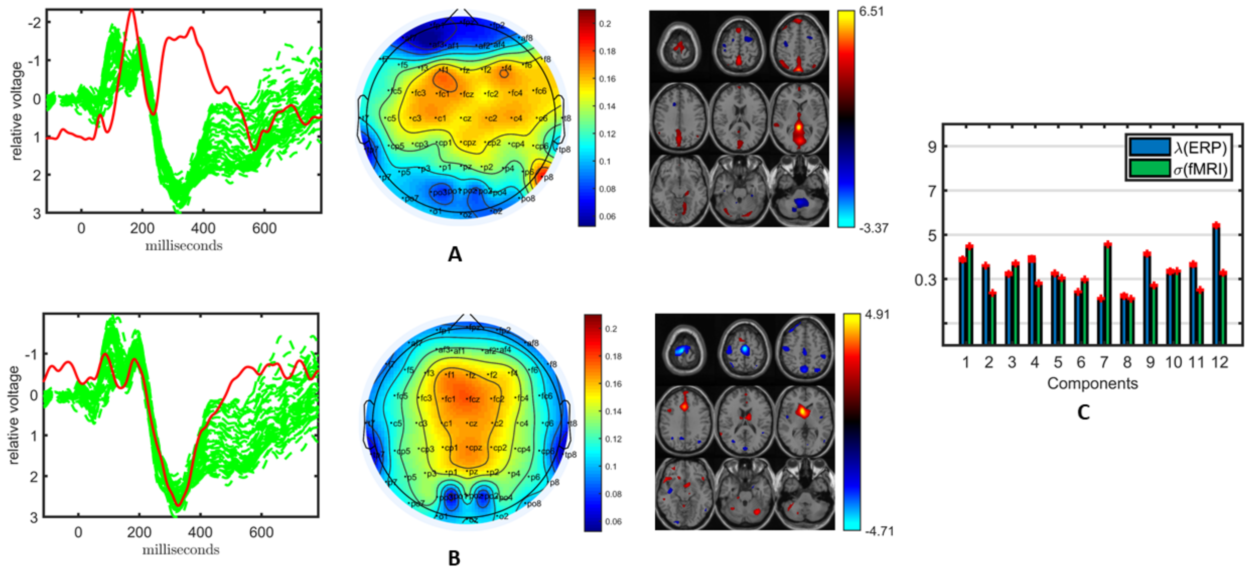


Figure 1: Temporal and spatial patterns from the statistically significant components of the ACMTF model of the EEG tensor with 62 electrodes and fMRI data, with $R = 12$. **(A)** Component 5 represents the P2-N2 as well as the P3 peak and is heavily weighted by the frontal and central electrodes in the EEG, while the fMRI shows increased activation of controls over patients in the sensorimotor cortex, **(B)** Component 12 refers to the P3 peak, heavily weighted by the central electrodes in the EEG and the fMRI indicates a decrease in activation of controls versus patients in some regions of the sensorimotor cortex, **(C)** Weights of the components in each data set. The corresponding $p$-values are $5.3 \times 10^{-3}$, $3.9 \times 10^{-4}$, respectively. Columns of the factor matrix in the *time* mode are in red while green plots show signals from individual electrodes averaged across all subjects.

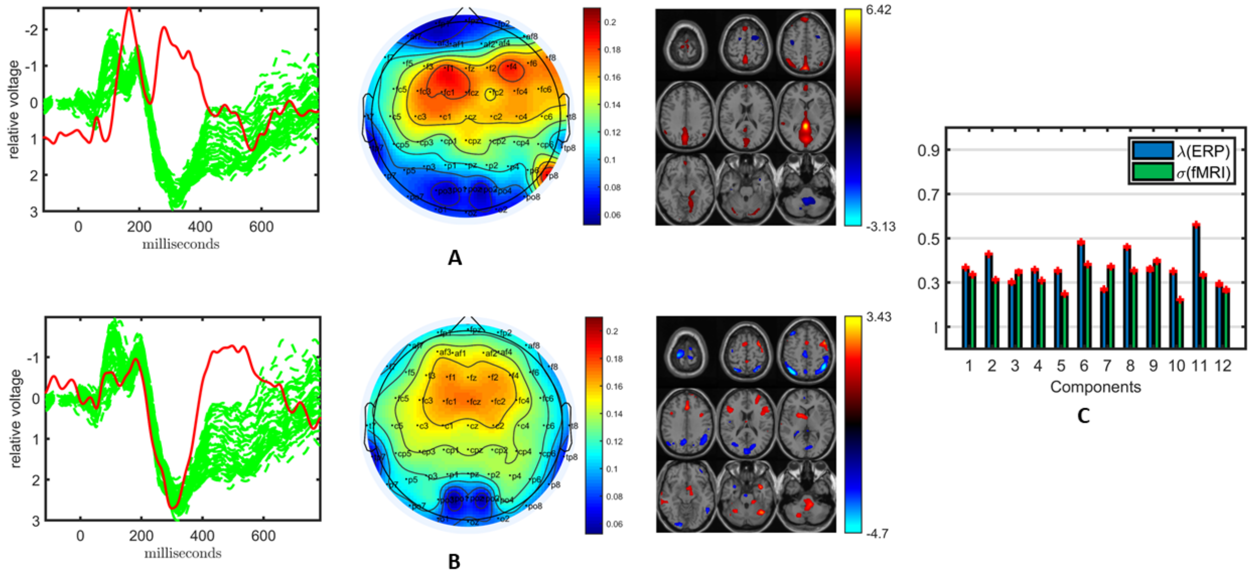


Figure 2: Temporal and spatial patterns from the statistically significant components of the ACMTF model of the EEG tensor with 62 electrodes and fMRI data, with $R = 12$, and with centering across the *subject* mode. **(A)** Component 9 refers to the P2-N2 as well as the P3 peak heavily weighted by the frontal and central electrodes in the EEG, while the fMRI shows increased activation of controls over patients in the parietal lobe, **(B)** Component 10 is related to the P3 peak and heavily weighted by the frontal electrodes in the EEG, while the fMRI indicates a decrease in activation of controls versus patients in the parietal lobe, **(C)** Weights of the components in each data set. The corresponding $p$-values are $3.8 \times 10^{-2}$, $4.8 \times 10^{-5}$, respectively. Columns of the factor matrix in the *time* mode are in red while green plots show signals from individual electrodes averaged across all subjects.

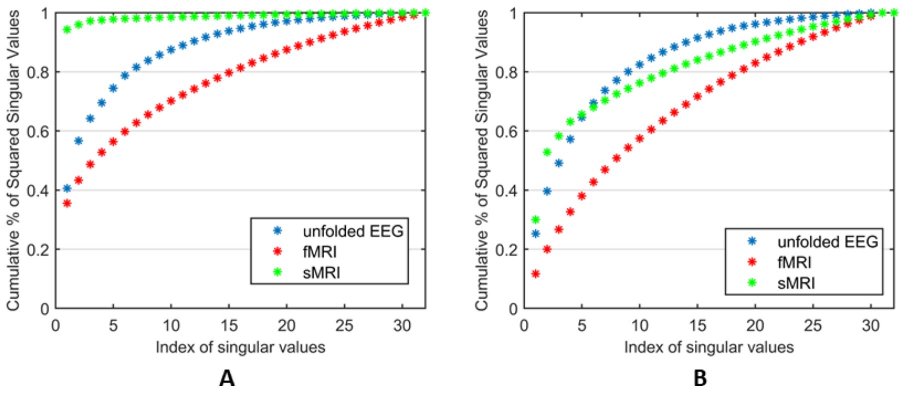


Figure 3: Cumulative percentage of the squared singular values for each data set. **(A)** No additional centering across the *subject* mode, **(B)** With additional centering across the *subject* mode. The EEG tensor is unfolded in the *subject* mode and arranged as a *subject* by *time-electrode* matrix before computing its singular value decomposition.

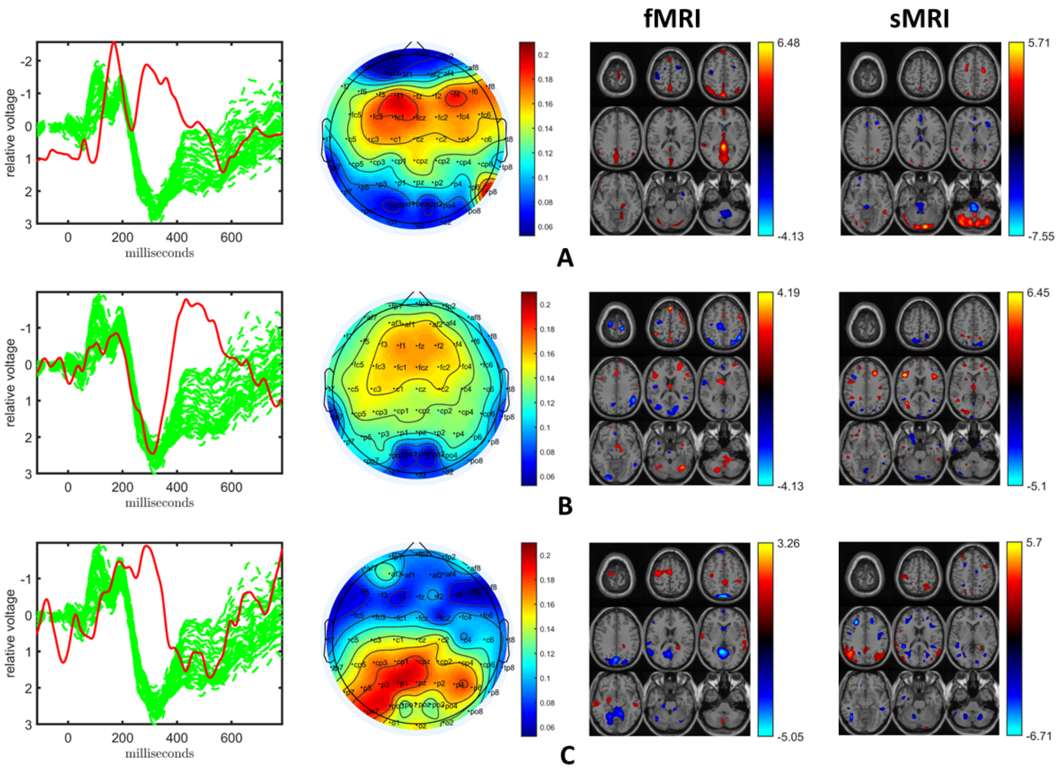


Figure 4: Temporal and spatial patterns from the statistically significant components of the ACMTF model of the EEG tensor with 62 electrodes, fMRI and sMRI data, with $R = 10$, and with centering across the *subject* mode. **(A)** Component 8 refers to the P2-N2 as well as the P3 peak heavily weighted by the frontal and central electrodes in the EEG, while the fMRI shows increased activation of controls over patients in the parietal lobe, and the sMRI shows increases in gray matter in controls over patients in the sections of the cerebellum, **(B)** Component 7 is related to the P3 peak and heavily weighted by the frontal electrodes in the EEG, while the fMRI indicates a decrease in activation of controls versus patients in the parts of the parietal lobe, and the sMRI shows increases in gray matter for controls over patients in some sections of the frontal lobe, **(C)** Component 5 refers to the N2 as well as P3 peaks, heavily weighted by the parietal electrodes in the EEG, while the fMRI shows increased activation of controls over patients in the sensorimotor cortex and a decrease in activation of controls versus patients in the occipital lobe. The corresponding sMRI component highlights changes to gray matter concentration throughout the frontal and parietal lobes. The corresponding $p$-values are $5.2 \times 10^{-3}$, $1.3 \times 10^{-2}$, and $3.4 \times 10^{-2}$ respectively. Columns of the factor matrix in the *time* mode are in red while green plots show signals from individual electrodes averaged across all subjects.

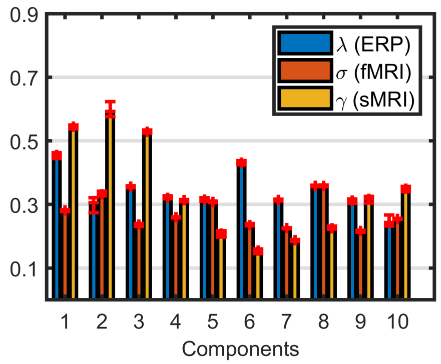


Figure 5: Weights of the components in EEG, fMRI and sMRI extracted by the ACMTF model of the EEG tensor with 62 electrodes, fMRI and sMRI data, with $R = 10$. Note that for these results there has been additional centering across the *subject* mode.
